# Risk‑Based Prediction of Novel AMR Variants Using Protein Language Models

**DOI:** 10.1101/2025.09.12.672331

**Authors:** Jennifer J Wood, Stephanie Portelli, David B Ascher, Nicholas Furnham

**Affiliations:** Department of Infection Biology, London School of Hygiene and Tropical Medicine, Keppel Street, London, WC1E 7HT UK; School of Chemistry and Molecular Biosciences, The University of Queensland, Saint Lucia Campus, Saint Lucia, Queensland, Australia

## Abstract

Antimicrobial resistance (AMR) is among the most pressing global health threats of the 21st century, with the potential to thrust modern medicine back into a pre-antibiotic era. Resistance can arise through diverse mechanisms, including genomic mutations that prevent antibiotics from reaching or acting on their targets. To limit the spread of AMR, surveillance systems must detect both known and emerging resistance markers. Here we present AMRscope, a model trained on ESM2 protein language model embeddings of single mutations for prediction of resistance likelihood, combined with a rigorous evaluation framework. This tool is applied across antibiotic-interacting proteins of different bacterial species, including WHO priority pathogens, such as rifampicin-resistant *M. tuberculosis* and carbapenem-resistant *P. Aeruginosa*. Performance on random splits achieves a competitive accuracy, F1 and MCC of 0.88, 0.87 and 0.75, respectively, while additional splitting strategies demonstrate transfer of predictive power to unseen organisms or genes. Moreover, *in silico* deep mutational scanning and structural mapping across these targets reveals the tool can recover known resistance-associated regions and highlight new candidates. The risk-based outputs complement database matching and resistance element detection tools, providing clinicians and public health agencies with an interpretable and scalable system for AMR surveillance and proactive response.

## Introduction

The emergence and spread of Antimicrobial resistance (AMR) is fast becoming a global public health crisis, with drug-resistant infections already causing an estimated 700,000 deaths annually, projected to rise to a total of 39 million over the next 25 years [1]. Consequently, the ability to promptly detect emerging resistance and understand the underlying mechanisms is vital for guiding public-health strategies and effective treatments. Pathogenic bacteria can be intrinsically resistant to an antibiotic, or resistance can be acquired through horizontal gene transfer or genomic point mutations.

Resistance arises either by altering a drug’s action or its access to the target, for instance through increased efflux pump expression, modified membrane permeability, antibiotic inactivation, disrupted prodrug activation, or direct target modification [2]. Given this dynamic and multifactorial nature of AMR, it must be proactively addressed through constant surveillance, antibiotic stewardship and development of novel treatments and predictive tools [3].

There are many existing computational methods for prediction of AMR phenotypes falling under two main methodologies: rule-based database mapping or machine learning approaches [4, 5]. Rule-based methods use matching to catalogues of known variants, and thus excel at prediction of previously seen resistance determinants but are constrained by prior knowledge and database quality, performing poorly on prediction of novel resistance determinants [6, 7]. ML based approaches offer more generalisability outside of known databases, with different input features and classification algorithms being used to identify underlying patterns in data, even in the absence of mechanistic understanding. Previous ML efforts have taken various forms, with models being trained on single or combinations of features, such as known SNPs [8], k-mer representations [9], physiochemical features [10] and evolutionary conservation [11]. While the use of hand-engineered features for model training can offer interpretability and mechanistic insight, such properties can be time consuming to extract and models are often gene-and species-specific, making them less scalable. In contrast, deep learning models such as those built on raw sequences [12] can be more generalisable and scalable but suffer from reduced transparency. There is a growing recognition that models must balance predictive performance with explainability, especially for applications in clinical or surveillance settings where actionable insights are required [13].

Recent advances in protein language models (PLMs), offer a promising foundation for striking this balance. Trained on millions of natural sequences, these models embed amino acid sequences into high-dimensional spaces that capture structural and evolutionary properties without the need for manual feature extraction [14-16]. The Evolutionary Scale Models (ESM) are one such series of transformer-based models, which have demonstrated strong performance in a range of mutation effect prediction tasks, particularly in human genomics [17]. For example, Varipred which uses ESM1b embeddings of mutant positions achieved performance comparable to, or exceeding, state-of-the-art tools in the most recent CAGI challenge [18]. However, this approach has not yet been applied extensively to AMR, where PLM-based models could be applied specifically to the problem of predicting resistance phenotypes from missense mutations in a pan-species context.

In this work, we present AMRscope, a predictive tool that estimates resistance risk from single amino acid substitutions in antibiotic target proteins. When applied across multi-species datasets, the model achieves competitive performance and shows transfer of predictive power to unseen organisms and genes, while *in silico* deep mutational scanning and structural mapping reveal recovery of known resistance hotspots alongside new candidates. Furthermore, AMRscope provides a configurable framework that unifies sequence preprocessing, embedding generation, flexible classifier training, and evaluation under diverse hold-out strategies. Together, these features position AMRscope as a scalable and interpretable tool for surveillance, leveraging protein language models against one of the most urgent challenges in infectious disease.

## Results

### Modelling of AMR mutations in M. tuberculosis

To develop baseline models for prediction of antibiotic resistance from missense variants, data from *Mycobacterium tuberculosis* was used, as genomic mutations are known to be a major driver of *M. tuberculosis* resistance [19]. Mutation data was shared by authors of tb-profiler [20] and filtered to produce a dataset of sensitive and resistant point mutations for genes with known resistance mutations - EmbB conferring ethambutol resistance [21], KatG for isoniazid resistance [22], PncA for pyrazinamide resistance [23], RpoB for rifampicin resistance [24], GidB for Streptomycin resistance [25] and Alr for cycloserine resistance [26]. For the initial modelling, a range of features were curated representing biophysical properties, evolutionary conservation, stability, and residue specific properties for the mutant position (summarised in table 1). In addition, ligand binding features were extracted using either existing crystal complex structures for EmbB-EMB (pdb id 7BVF) [27] and RpoB RNAP β-rifampicin (pdb id 5UHC) [28] or by generating docked complexes using auto dock vina [29] for PncA-pyrazinamide, KatG-isoniazid, Alr-cycloserine, GidB-streptomycin pairings.

**Table 1.**
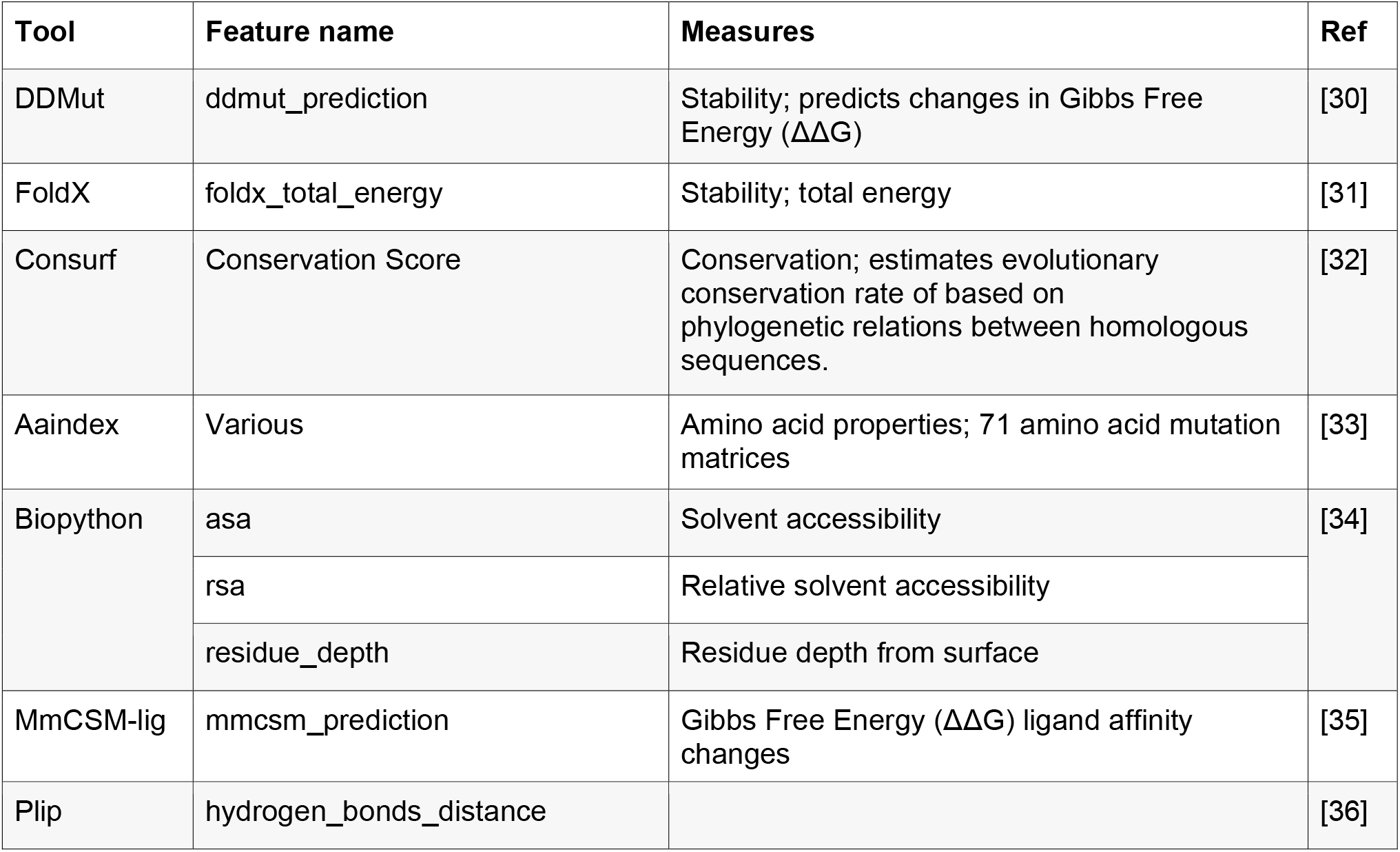
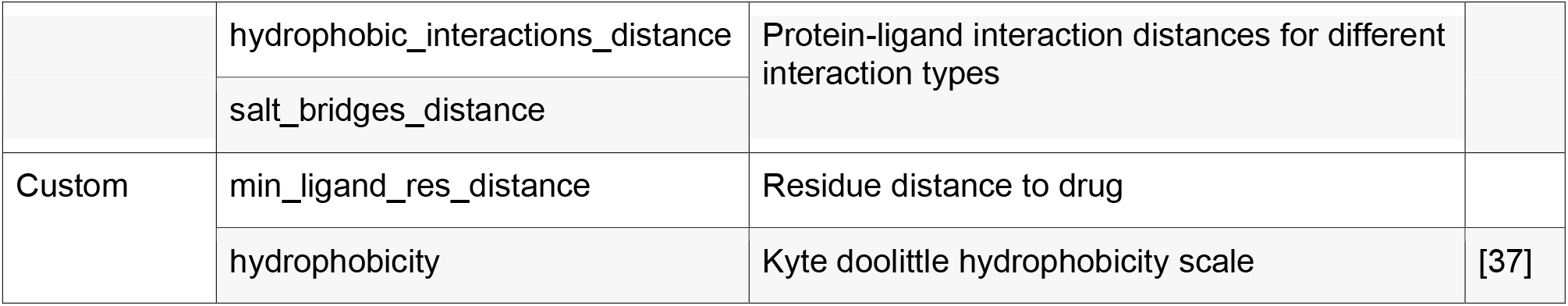
Feature based model inputs. Table summarising the features used for model training prior to feature selection and the corresponding tools.

Comparative modelling was performed using logistic regression (LR), gradient boosted classifier (GBC), balanced random forest (BRF) and gaussian process (GP) models, with automatic feature selection to determine the best set of predictive features. In this approach we equate drug resistance to a binary classification problem, with 0 (negative class)/1 (positive class) labels for antibiotic sensitivity and resistance respectively. Modelling was carried out individually for genes with sufficient datapoints (EmbB, KatG, PncA, RpoB), in addition to a combined model across all six genes. Data was randomly split into train and test sets while preserving the ratio of resistant and sensitive datapoints. Throughout this study we report accuracy (or balanced accuracy), F1-score, Matthews correlation coefficient (MCC), precision and recall, however we focus on F1 score and MCC as our primary metrics of interest. F1 score considers the balance of precision and recall with values from 0-1, where in general a score above 0.6 is considered acceptable performance. MCC considers all possible outcomes of a confusion matrix, ranging from -1 to +1, with a value above 0.3 considered moderate and above 0.5 considered strong correlation.

From the summary of test set performance statistics (table 2) we see that, while in some cases overall performance is better with larger and more balanced (resistant : sensitive) datasets, this is not always true, and thus it is also gene dependent. For example, the model of RpoB resistance to rifampicin performs well, with an F1 score of 0.71, an MCC of 0.69 and recall of 0.89, in line with previously published tools, where resistance is known to be strongly linked to mutations near the rifampicin binding pocket [11]. Conversely the KatG model performs poorly, with an F1 of 0.27, MCC of 0.15 and recall of 0.52, despite there being similar number of datapoints to the RpoB resistance model. Furthermore, the balanced random forest performs best across most of the datasets, likely compensating for some of the imbalance in resistant to sensitive datapoint ratios.

**Table 2.** AMR prediction performance on the *M. tuberculosis* dataset. Table summarising the number of positive and negative datapoints in each dataset and the test-set performance metrics for the best performing feature-based model.

| Dataset | N sensitive datapoints | N resistant datapoints | % Resistant | Best Model | Balanced Accuracy | F1 Score | MCC | Precision | Recall |
| --- | --- | --- | --- | --- | --- | --- | --- | --- | --- |
| Combined | 4263 | 539 | 11.22 | BRF | 0.81 | 0.56 | 0.51 | 0.44 | 0.77 |
| PncA | 230 | 252 | 52.28 | LR | 0.77 | **0.78** | 0.55 | **0.74** | 0.83 |
| RpoB | 1080 | 114 | 9.55 | BRF | **0.91** | 0.71 | **0.69** | 0.60 | **0.89** |
| **KatG** | 985 | 102 | 9.38 | BRF | 0.61 | 0.27 | 0.15 | 0.18 | 0.52 |
| **EmbB** | 1092 | 61 | 5.29 | BRF | 0.76 | 0.34 | 0.33 | 0.23 | 0.18 |
| **GidB** | 602 | 8 | 1.31 | - | - | - | - | - | - |
| **Alr** | 274 | 2 | 0.72 | - | - | - | - | - | - |

The best combined model, containing information from all our gene sets, performs reasonably with a balanced accuracy of 0.81, F1 of 0.56, MCC of 0.51 and recall of 0.77. To confirm that this performance is not sensitive to dataset splits we also performed 5-fold cross validation with assessment on a validation subset, which gives consistent performance, indicating the BRF model is robust (Figure 1a). Furthermore, to ensure this performance was not driven wholly by a single gene set, prediction classes of our test set (True or False, Negative and Positive) were visualised (Figure 1b). Here we see that while there is a difference in performance, all four individual gene predictions have some true negatives and some true positives.

**Figure 1.**
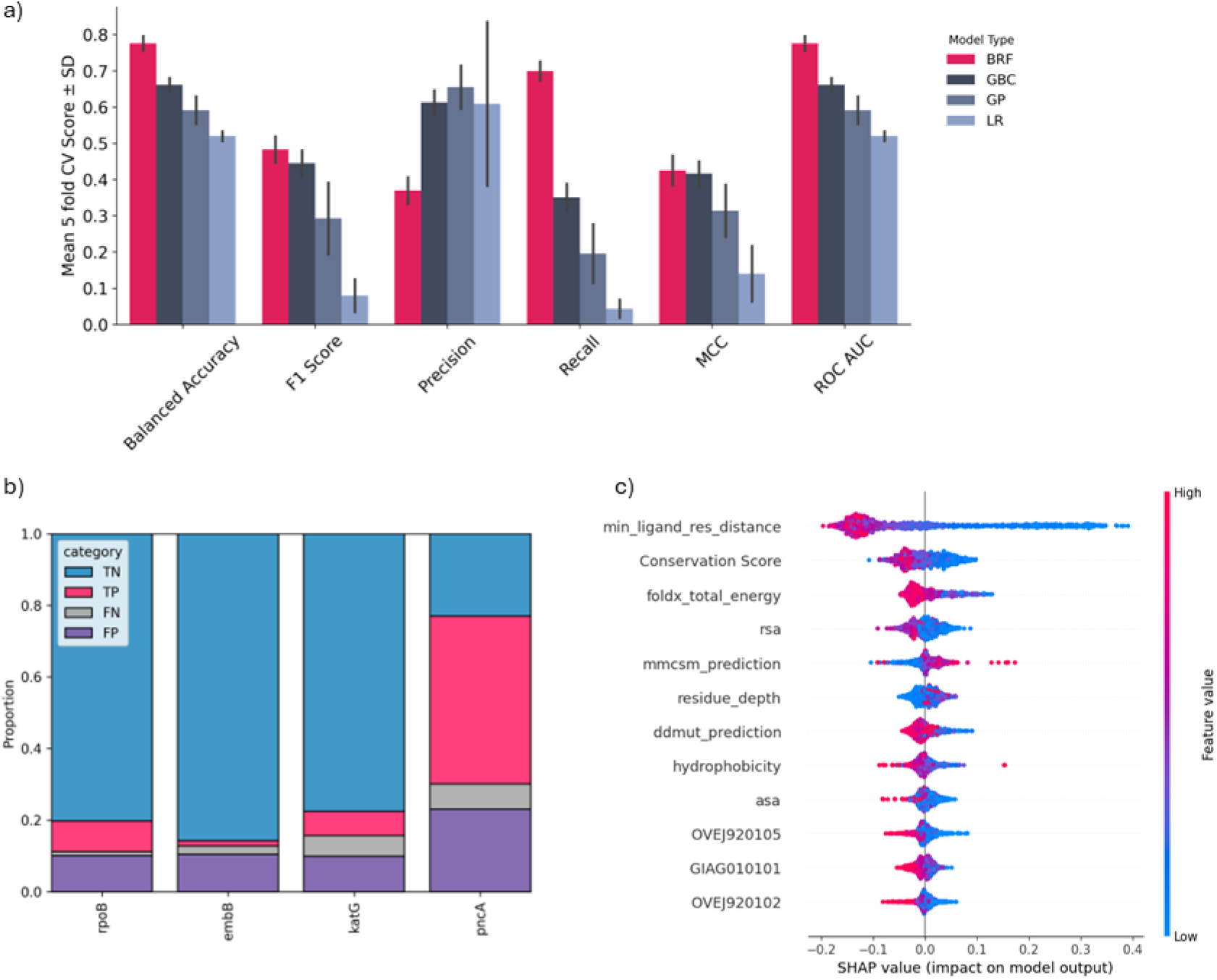
Prediction of AMR on a mixed gene *M. tuberculosis* dataset. a) Test set performance of four different feature-based binary classification models with 5-fold cross-validation (Balanced Random Forest (BRF), Gradient Boosting Classifier (GBC) Gaussian Process (GP) and Logistic Regression). b) Classification categories of True/False, Positive/Negative for the best combined model (BRF) predictions split by the four major represented genes. c) SHAP feature importance from the best combined model.

To generate additional insights from this model, the individual feature impacts were evaluated using feature importance and SHAP summary plots. Figure 1c illustrates the 12 automatically selected features and their impact upon predictions on the test set from our combined model (Figure S1 for single gene SHAP plots). The descriptions of most of these features can be found in Table 1, with the remaining three OVEJ920105, OVEJ920102, GIAG010101, being individual substitution matrices from aa index, which represent environment-specific amino acid substitution matrix for inaccessible residues, environment-specific amino acid substitution matrix for alpha residues, and residue substitutions matrix from thermo/mesophilic to psychrophilic enzymes respectively [33]. Considering these properties, the distance between ligand and residue has the greatest impact, where as expected, mutations closest to the antibiotic binding site have a high positive impact on the prediction of resistance. The remaining selected features cover conservation, stability prediction, position within the molecule and amino acid properties.

Following the evaluation of our baseline models, a workflow was developed to leverage embeddings from the ESM-2 PLM [17] for single-residue mutation classification. Given the small size of our dataset, we used the pretrained (frozen) ESM2 weights to generate the embeddings for classification, without additional training of the ESM2 model itself. For each variant, the wild-type and mutant sequences were tokenized and embedded using the pretrained PLM, and the final-layer vectors at the mutated position were extracted and concatenated into a single feature vector and the log-likelihood ratios appended as the final element. This strategy preserves both the mutation-specific signal and its surrounding sequence context. Additional curated features could also be appended to the feature vector to determine if supplying the model with more information could further improve performance, and these final vectors were used to train a single layer MLP classifier.

This pipeline was used to again evaluate models of individual EmbB, KatG, PncA, RpoB genes, in addition to a combined model across all resistance genes, with dataset splits matching the previous section to allow direct comparison. Test set results in Table 3 illustrate that, like feature-based models, performance is related to the number of training datapoints and the specific gene. This PLM embedding approach results in a better F1 score and MCC on the combined data set and with RpoB but performs even more poorly on the KatG and EmbB datasets. None of the results were significantly improved by taking an ensemble of predictions from the two modelling approaches, or by training the classification layer on a concatenated vector of embeddings and hand curated features.

**Table 3.** AMR prediction performance using PLM embeddings on the *M. tuberculosis* dataset. Performance metrics of the PLM based model for each dataset averaged with 5-fold cross validation.

| Dataset | Test Performance |  |  |  |  |
| --- | --- | --- | --- | --- | --- |
|  | Accuracy | F1 Score | MCC | Precision | Recall |
| Combined | 0.92 | 0.66 | 0.61 | 0.66 | 0.66 |
| PncA | 0.57 | 0.68 | 0.19 | 0.54 | **0.94** |
| RpoB | **0.96** | **0.84** | **0.81** | **0.84** | 0.83 |
| KatG | 0.89 | 0.09 | 0.12 | 0.35 | 0.05 |
| EmbB | 0.92 | 0.27 | 0.26 | 0.44 | 0.20 |

### Collation of a multi-species AMR mutation benchmark dataset

Given the improved performance of the PLM embedding based models on larger datasets, we sought to expand our training set to determine if performance could improve further and if our model could perform across different species. To achieve this, data on AMR mutations was collected from a variety of sources and combined with our *M. tuberculosis* data, including the Comprehensive Antibiotic Resistance Database (CARD) [38], AMR finder [39], pathogen watch [40, 41] and pointfinder databases [42]. Resistance mutations were downloaded from these sources and filtered for missense mutations, any duplicate mutations were dropped to leave only unique mutations and these sequences were filtered to a set which are known or suspected to be direct antibiotic binders (appendix table S1).

Unlike the *M. tuberculosis* dataset, the aforementioned databases do not contain sensitive mutation data, as is required for classifier model training, and thus one needed to be separately constructed. To avoid the use of purely synthetic negative data, an additional pipeline was developed to extract mutations from the ‘AllTheBacteria’ (v0.2) [43], a database containing all WGS isolates from the bacterial International Nucleotide Sequence Database Collaboration data which were uniformly assembled, QC-ed and annotated with results from AMRFinder (data collation summarised Appendix figure S2). This final combined dataset contains 3907 unique datapoints, with 2123 sensitive and 1784 resistant mutations, from 36 genes across 46 species, including WHO priority pathogens. It should be noted there is a large range in numbers represented from each group but, as our model is focused on the individual protein level, it was ensured that every gene has representation in both the sensitive and resistant class (appendix table S2).

To assess the ability of the model to generalise out of the training distribution, several different data splitting strategies were implemented in addition to the initial random stratified split. This includes splitting by gene, organism, or antibiotic groups, where there is no overlap between the train and test set for the indicated group. Homology based splits were performed based on splitting by both CATH domains [44] and cd-hit clusters (threshold = 0.4) [45]. Each training set was subjected to further 5-fold cross validation splits to assess the robustness of the model.

### Modelling of AMR mutations across genes and species

To provide a comprehensive evaluation of the new multi-species dataset, the modelling workflow was extended to produce embeddings using ESM-1b, ESM-2 or SaProt, where SaProt is used to incorporate local structure information into the modelling process [17, 46] and training was performed with different classification heads (Figure 2). Again, we specifically maintained a binary classification approach, as there were not enough datapoints per antibiotic class to support multi-class classification. Figure 3a illustrates performance on the random test set with 5-fold cross-validation, where we compare the best performing classifier with ESM-1b and SaProt, to performance of all classifiers with the best performing ESM2 embeddings. Considering F1 score and MCC as primary metrics, we find the ESM2 with a single layer MLP performed best, with F1 of 0.87 and MCC of 0.76, which is an improvement on our combined dataset using the imbalanced *M. tuberculosis* only dataset. The associated ROC and Precision recall curves of all folds also show consistent high ROC-AUC and PR-AUC, demonstrating stable and effective discrimination between resistance and sensitive mutations (figure 3b).

**Figure 2.**
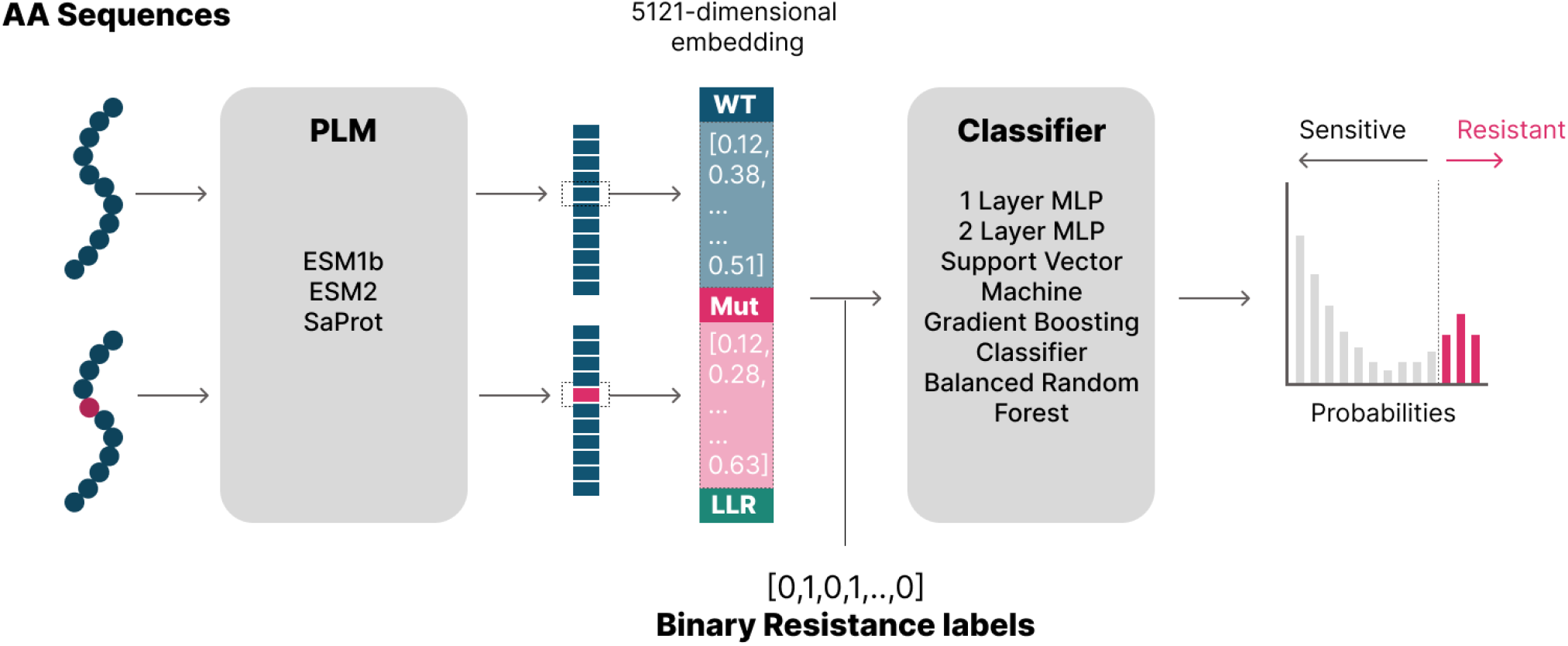
PLM-based AMR classification pipeline. Embeddings are generated for wild type (WT) and single mutant sequences and feature vectors extracted from the final model layer at the mutant position. WT and mutant embeddings are concatenated into a single vector and the log-likelihood ratios appended. These vectors were used to train a binary classification layer with resistant/sensitive (1/0) target labels.

**Figure 3.**
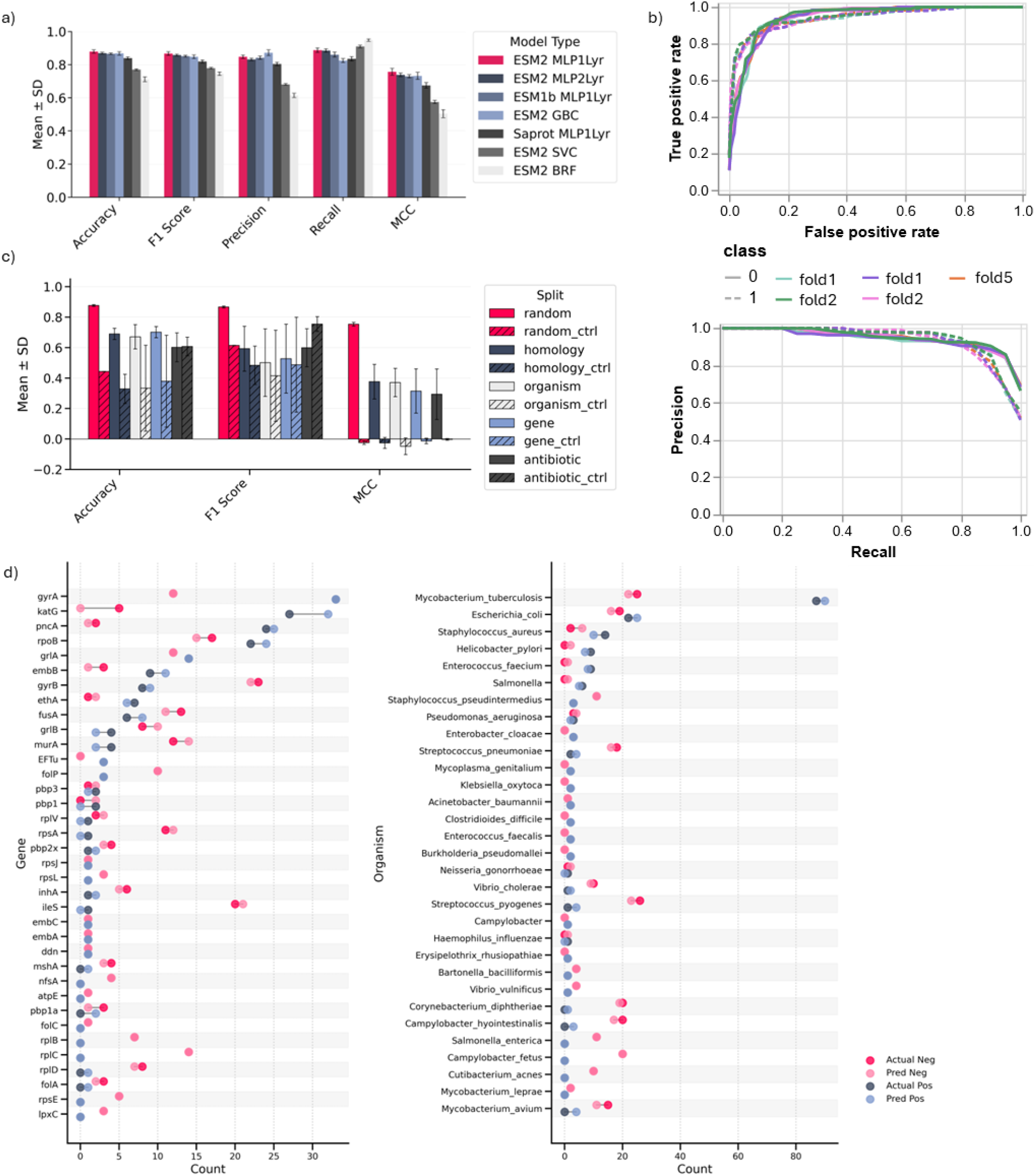
Modelling performance and comparison on a multi-species, multi-gene dataset. a) Test set performance comparison using different embeddings (ESM2, ESM1b, SaProt) and classifiers (1-layer MLP, 2-layer MLP, GBC, SVC, BRF) with 5-fold cross validation. b) ROC and Precision-Recall curves for five folds of ESM2 MLP 1layer model c) Performance of ESM2 1-layer MLP model retrained on data with different hold out splits to evaluate on unseen antibiotics, genes, organisms or homology groupings. Five holdouts were assessed for each split and averaged, and a baseline is obtained by randomising training labels to provide a no signal control. d) Dumbbell plots of actual/predicted - positives/negatives, split by gene or organism.

Given our model covers many different species and genes, dumbbell plots of actual and predicted positives and negatives split by these groups were also generated for representative predictions on the test set (Figure 3d). Here we see good correspondence between predictions and ground truth for some subsets, for example GyrA and *M. tuberculosis*, and poorer for other subsets such as KatG, which again seems intractable to our modelling approach.

To assess the ability of the model to generalise to out-of-distribution mutations, the selected best architecture was retrained and tested using five distinct hold-out strategies, each with five non-overlapping test groups (summarized in Table S3). Performance metrics were averaged across the five folds for each strategy. Figure 3c displays bar plots of the mean ± standard deviation for Accuracy, F1 Score, and MCC, grouped by data split. Baseline control models were generated by randomly shuffling the resistant/sensitive labels during training, to illustrate performance based on no true signal, and are shown as hashed bars vs the solid bars correspond to models trained on real labels.

Using F1 and MCC as primary metrics, the model performed well on the random split (F1 = 0.87, MCC = 0.75), with more modest but still meaningful performance across biologically structured splits: homology (0.59/0.37), organism (0.50/0.37), gene (0.53/0.31), and antibiotic (0.60/0.29). Importantly, the model consistently outperformed its corresponding controls in MCC, which accounts for all elements of the confusion matrix and is robust to class imbalance. This gap highlights that the model is learning genuine biological signal rather than overfitting to noise. However, generalization to more distantly related unseen data is clearly influenced by the difficulty of each specific hold-out set.

To our knowledge, there is currently no benchmark dataset or like-for-like model which specifically uses single mutation-based inputs for pan-AMR predictions to compare with our approach. Therefore, we used the following three general models to provide an indication as to the utility of our model vs alternative generally available tools for this problem space that could be utilized for this prediction; 1) Zeroshot-ESM2 scoring (delta logP score, threshold = 0), 2) Zeroshot-Prot-T5 scoring (cosine similarity, threshold = 0.99) and 3) VESPA-G [47, 48]. In this case the comparison with ESM2 highlights the uplift from fine-tuning, while ProtT5 compares the performance of an alternative unsupervised encoder/decoder protein language model. Currently, VESPA-G is the top performing sequence-only variant effect predictor on protein gym, a collection of benchmarks aiming at comparing the ability of models to predict the effects of protein mutations [49]. The VESPA-G model also uses ESM2 and a supervised head but trained across diverse mutational data sources, providing a broader, but supervised comparator. Table 4 provides metrics for mutations for a single hold out set for each scenario, illustrating that our AMR specific model performance improves upon more general models across the different test sets. Although these comparisons are informative in terms of fine-tuning uplift for our data, we emphasize that they are not true benchmarks in the absence of an independent standardized test dataset for pan-AMR missense prediction and reference models.

**Table 4.** Summary of performance metrics of AMRscope vs selected models on a single test set per hold-out type.

| Split | Model | Accuracy | F1 | MCC | Precision | Recall |
| --- | --- | --- | --- | --- | --- | --- |
| random | AMRscope | 0.88 | 0.87 | 0.76 | 0.82 | 0.93 |
|  | ESM2-ZeroShot | 0.52 | 0.64 | 0.22 | 0.48 | 0.96 |
|  | ProtT5-ZeroShot | 0.57 | 0.63 | 0.23 | 0.51 | 0.83 |
|  | VESPAG | 0.32 | 0.43 | -0.35 | 0.33 | 0.59 |
| organism | AMRscope | 0.73 | 0.37 | 0.41 | 0.23 | 1.00 |
|  | ESM2-ZeroShot | 0.19 | 0.13 | 0.10 | 0.07 | 1 |
|  | ProtT5-ZeroShot | 0.32 | 0.14 | 0.08 | 0.07 | 0.87 |
|  | VESPAG | 0.12 | 0.09 | -0.23 | 0.05 | 0.64 |
| homology | AMRscope | 0.67 | 0.64 | 0.39 | 0.62 | 0.72 |
|  | ESM2-ZeroShot | 0.48 | 0.58 | 0.17 | 0.42 | 0.93 |
|  | ProtT5-ZeroShot | 0.52 | 0.56 | 0.14 | 0.44 | 0.77 |
|  | VESPAG | 0.31 | 0.42 | -0.32 | 0.31 | 0.64 |
| gene | AMRscope | 0.73 | 0.42 | 0.41 | 0.28 | 0.91 |
|  | ESM2-ZeroShot | 0.27 | 0.21 | 0.13 | 0.12 | 0.98 |
|  | ProtT5-ZeroShot | 0.39 | 0.23 | 0.16 | 0.13 | 0.90 |
|  | VESPAG | 0.10 | 0.04 | -0.57 | 0.02 | 0.22 |
| antibiotic | AMRscope | 0.55 | 0.50 | 0.30 | 0.91 | 0.35 |
|  | ESM2-ZeroShot | 0.71 | 0.82 | 0.31 | 0.69 | 0.99 |
|  | ProtT5-ZeroShot | 0.66 | 0.77 | 0.16 | 0.69 | 0.85 |
|  | VESPAG | 0.30 | 0.42 | -0.45 | 0.36 | 0.50 |

### Interrogation of embeddings, model reliability and predictions

To understand the structure of the PLM-derived embeddings and assess the presence of underlying patterns, we applied unsupervised dimensionality reduction using t-distributed Stochastic Neighbour Embedding (t-SNE) and Principal Component Analysis (PCA). t-SNE was used to project the input vectors into two dimensions, with ground truth labels overlaid (Figure 4) and separately, PCA was performed with 50 components on the same vectors and results were visualised as heat-maps grouped by organism, gene, wild-type and mutant amino acids. The t-SNE projection exhibited partial separation between positive and negative classes, indicating that some degree of linear separability exists in the latent representation space. The PCA heat-maps also revealed some structure within specific subgroups, particularly in the higher principal components. While these analyses are primarily qualitative, they reveal structure in the embeddings that correspond to biological groupings, even though such metadata are not explicitly provided. Notably, no single metadata category explains the full structure, suggesting a combination of latent features spanning multiple biological axes, including other learned PLM features such as evolutionary and physiochemical properties, can inform the classification layer.

**Figure 4.**
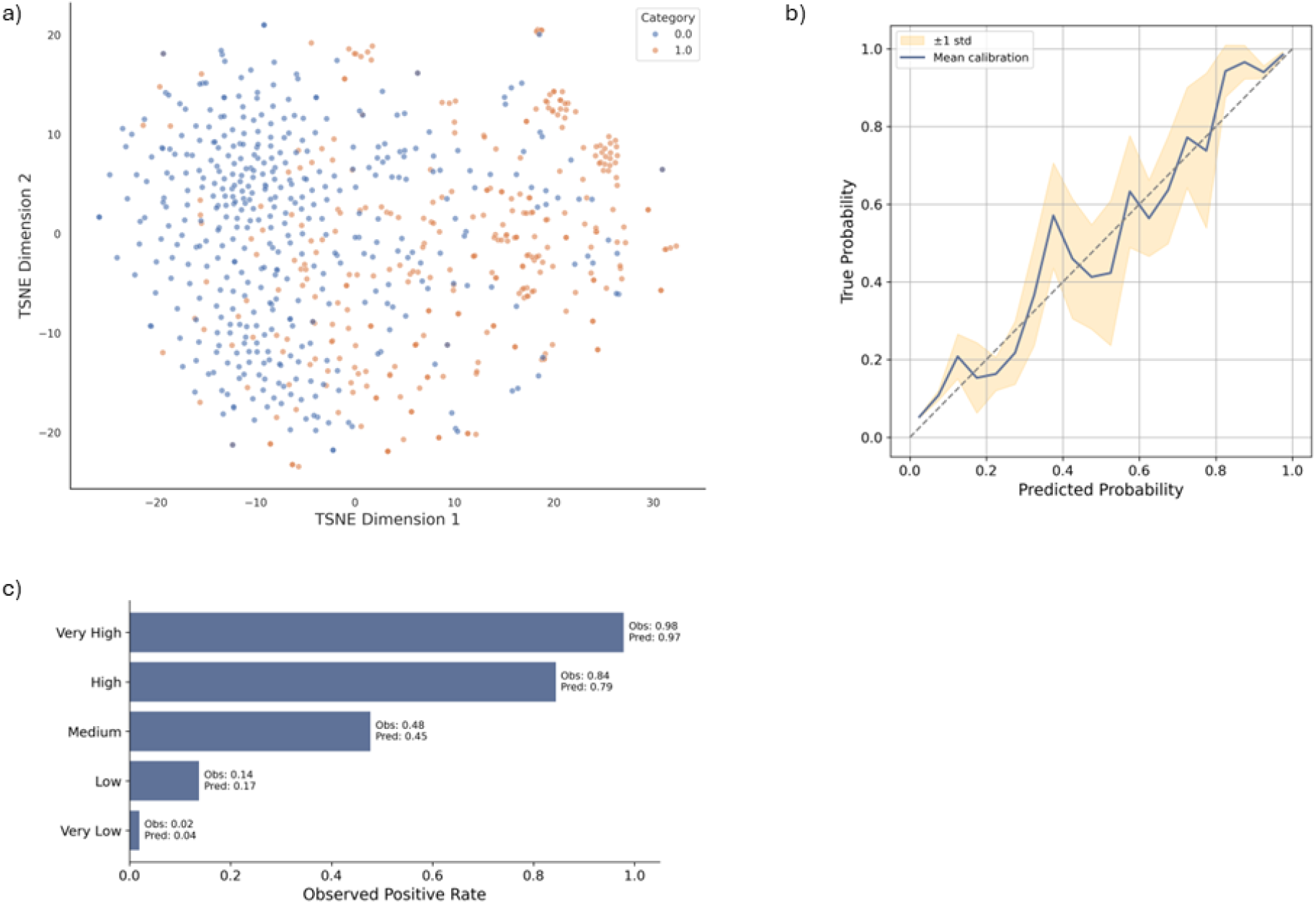
Interrogation of embeddings and reliability of predictions. a) t-SNE plot of reduced embeddings with ground truth label colour (0= sensitive / 1= resistant) for a representative validation set. b) Test set calibration curve of average predicted probabilities vs observed positive frequency for equal width bins. c) Definition of risk categories based upon expected probabilities.

To analyse how well the predicted probabilities reflect true outcomes, complementary to the evaluation of binary outcomes, we computed a calibration curve using the test set results (Figure 4c). This plot is constructed using the raw model output values, where 0 is lowest possible probability of a mutation causing resistance and 1 is the highest possible probability. Here we see the resulting calibration curve tracks along the diagonal, suggesting that the model is reasonably well-calibrated overall. Some deviation is observed in the mid-probability range, which is further supported by a Brier score of 0.09, indicating good but not perfect calibration. Taken together, these results suggest that the model’s probabilistic outputs are interpretable and can be used with some degree of confidence.

Given our calibration curve provides evidence for reliability of our probabilities, we defined five risk categories (very low, low, medium, high and very high) to provide granular, interpretable outputs alongside binary predictions. To derive the risk categories, we divided the predicted probabilities from all folds of all sets (train, validation, and test) into fixed intervals and computed the actual proportion of true positives observed in each bin. For example, in the very low (0.00–0.10) bin, the model’s average predicted probability was 0.04, of which 2% of those predictions were positive. At the other extreme, the very high (0.90–1.00) bin had an average predicted probability of 0.97 and an observed positive rate of 98% (Figure 4c). Again, the alignment between predicted and actual positive rates supports the model calibration, meaning there is no need for post-hoc alteration of bin definitions.

Finally, to interrogate the model behaviour per residue across specific genes, we conducted *in silico* deep mutational scanning (DMS) for specific targets, generating predictions for every possible single amino acid substitution across the protein. Illustrative genes covering different species and antibiotic resistances were selected for prediction, including three from the WHO priority pathogens list [50]; Rifampicin resistant *M. tuberculosis* (critical priority), carbapenem resistant *Pseudomonas Aeruginosa* (high priority), fluoroquinolone resistant *Salmonella* (high priority) and fluoroquinolone resistant *Escherichia coli*. To incorporate maximal data into the final model, which is used purely for this predictive inference and not model evaluation, AMRscope was retrained to include the test set data while retaining the same validation set for early stopping, which determines the optimal number of training iterations (epochs) based upon plateauing of the validation loss. Figure 5 illustrates results for a) *M. tuberculosis* RpoB, b*) E. coli* GyrB, c) *Salmonella* GrlA, and d) *P. Aeruginosa* PBP3, including heat map of predicted probabilities along the sequence length and average position probabilities on AlphaFold structures, in addition to a histogram of mutation risk categories. Datapoints for each of these genes present in our dataset are also summarized for reference in table S4.

**Figure 5.**
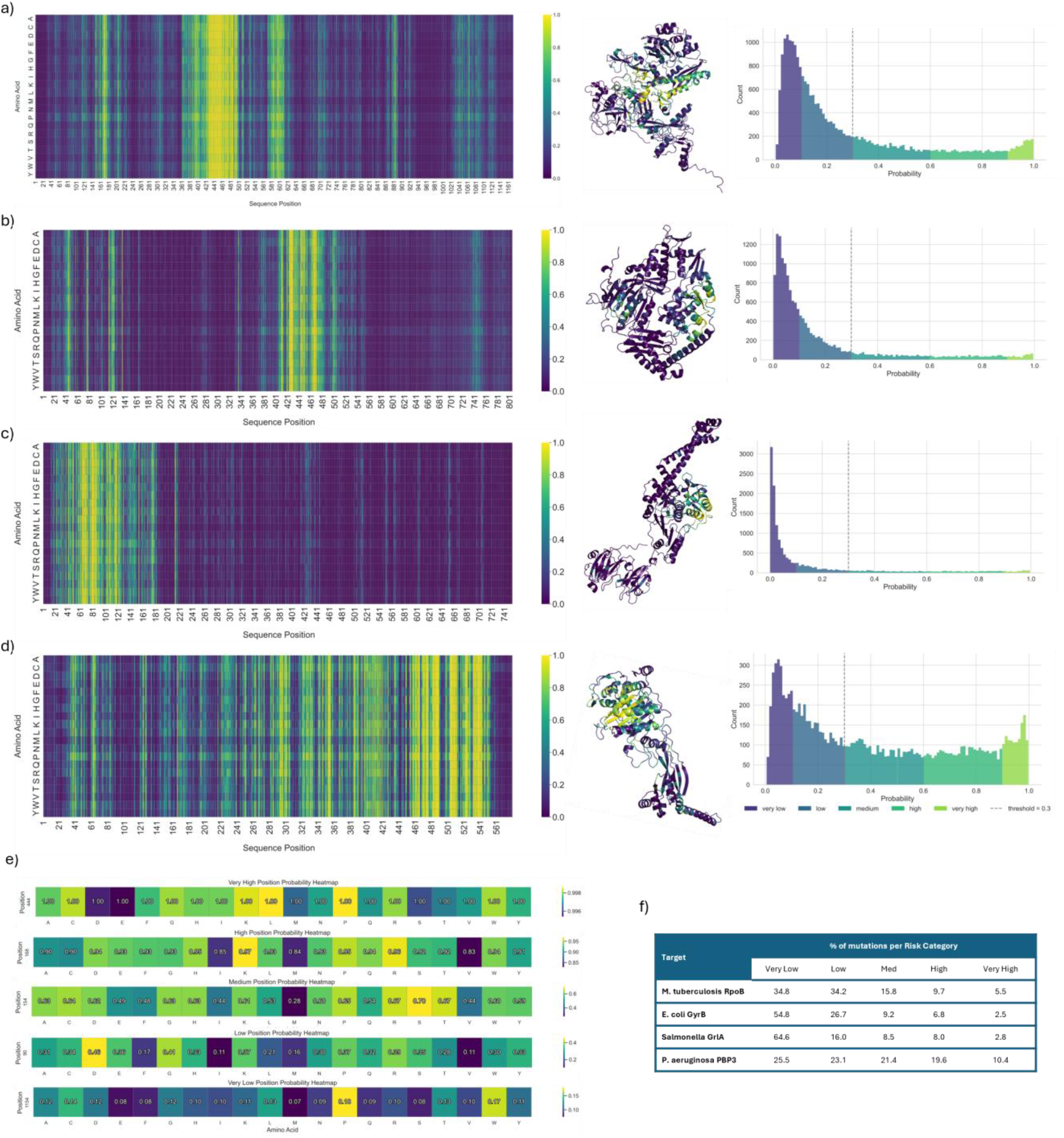
In Silico DMS prediction of a selection of high priority targets. Heatmaps of model predictions for every possible amino acid with averaged values mapped onto Alphafold structures and histogram of risk scores for a) *M. tuberculosis* RpoB b) *E. coli* GyrB c) *Salmonella* GrlA d) *P. aeruginosa* Pbp3. e) Example of amino acid probabilities for different risk categories of *M. tuberculosis* RpoB. f) Summary of percentage of mutations within each risk category.

For *M. tuberculosis* RpoB we see that the model is capable of highlighting both the known rifampicin resistance determining region (RDDR) around amino acids 426–452 [51] and resistance mutations outside of this region, such as the highly resistant V170F substitution [52]. Considering *E. coli* GyrB, we again see that *in silico* DMS highlights the known Quinolone Resistance-Determining Region (QRDR) in the window of 426–464 [53], even though there are only two quinolone resistant *E-coli* Gyrb mutations in our original dataset (D426N, K447E). This is likely reinforced by the equivalent resistant mutations in other species (such as *Salmonella* and *Clostridioides difficile*), since ESM2 embeddings are able to capture sequence context around the mutation. In the case of *Salmonella* GrlA, there is a known Quinolone Resistance Determinant Regions (QRDR) spanning amino acids 64-103 [54] which has the highest prediction values, including assigning ‘very high’ values to mutations G72S and D79G, which are not present in our dataset but we found to be reported as resistant from our independent literature search [55]. Lastly, *P. Aeruginosa* PBP3 is composed of two domains, an N-terminal domain (residues ∼1–221) and a C-terminal transpeptidase domain (∼222–577), where carbapenem binding and resistance-associated mutations map to the transpeptidase domain, particularly around the conserved catalytic motifs and adjacent loops (∼460–540) [56, 57]. Looking at the probabilities mapped to the 3D structure, we see the highest values in this transpeptidase domain and although in this instance the model seems to overpredict resistance probability, these are highly concentrated around the appropriate protein space.

As previously noted, due to the limited number of resistance annotations per antibiotic, we approached this problem from a binary perspective rather than predicting the specific antibiotic to which a mutation confers resistance. Antibiotic associations are instead inferred from resistant mutations in our curated database and can be retrieved for any gene in the dataset via a simple command-line interface. For example, retrieving resistance-associated mutations in *M. tuberculosis* RpoB, *Salmonella* GrlA, and *P. aeruginosa* PBP3 correctly returns rifampicin, fluoroquinolones, and β-lactams, respectively. In the case of *E. coli* GyrB, both fluoroquinolones and aminocoumarins are returned, however, resistance to fluoroquinolones can be inferred as the most likely explanation of resistance given proximity and correlation with mutations in other species.

Using RpoB as an example to look in more detail at the amino acid level, one mutation was randomly selected from each risk category and the heat map plotted for all amino acids at that position, with the minimum and maximum set separately for each position (figure 5e). In this example, we see there is variation in probability across the amino acids in all categories, although this is less for very high (0.02) and very low (0.09) bins than for the wider, intermediate categories such as medium risk (0.32). Overall, however, it seems that positional information dominates model predictions over amino acid residue changes.

## Discussion

In this study we present AMRscope, a tool which uses single site embeddings from the ESM2 protein language model, coupled with a binary classifier, to accurately flag missense mutations associated with antimicrobial resistance (AMR) across diverse antibiotic interacting genes and bacterial species. As using binary predictions alone masks any underlying uncertainty, we also chose to group variants into risk groups, supporting practical use in triage settings for novel variants. Our *in silico* deep mutational scans (DMS) can recapitulate known resistance associated regions and suggest novel candidate variants, illustrating how this model can complement existing database matching and mobile element detection tools to indicate potential of emergent resistance.

Focusing first on *M. tuberculosis*, we find that classifiers trained on PLM embeddings perform comparably to hand-curated properties on larger datasets, although performance degrades in cases of extreme sparsity (where p ≫ n), suggesting that ESM2 embeddings can effectively capture this feature information. Considering the most important hand-curated features, several of our identified mutation factors align with expectations from previous studies, for example distance to drug, ligand affinity, conservation and solvent accessibility for RpoB-rifampicin and PncA-PZA [11, 58, 59]. However, for KatG, neither PLM embeddings nor curated features yield accurate predictions, suggesting some targets may require a different approach. This may be due to KatG’s role in prodrug activation rather than being the direct target of inhibition, and the fact that resistance arises through allosteric mutations affecting conformational dynamics as opposed to active-site disruption. For KatG, prediction may be inherently difficult without incorporating protein flexibility or dynamic information, suggesting future work may benefit from incorporation molecular dynamics [60].

We further show that a single model trained on multi-gene, multi-species data can learn generalizable patterns of AMR-associated sequence variation. This capacity to transfer knowledge across genes and pathogens is demonstrated through DMS predictions, which reveals resistance-enriched regions across different targets, aligning with known resistance-determining motifs, even in cases where few labelled examples were seen during training for that specific protein. DMS predictions also assign high resistance probabilities to several mutations which are not present in the training data but supported by literature evidence, suggesting the model can flag novel mutations if and when they arise. We additionally mapped these residue-level predictions onto AlphaFold structures to aid spatial interpretation. While some overprediction is observed, likely due to the model’s limited ability to effectively penalize biophysically implausible mutations, the spatial clustering of high-risk predictions around active sites or known resistance hotspots further supports the biological plausibility of these outputs.

While the specific amino acid substitution contributes to predictions, we find that positional context is the primary driver of model output. This likely reflects the sparsity of data in terms of both breadth and depth of coverage at each position, which limits the model’s ability to fully learn amino acid substitution effects. It is noted that some of the amino acids with the highest predicted probability of resistance represent unlikely substitutions from a protein engineering perspective, for example substitution with biophysically unfavourable proline in the very high and high categories, which may reflect the fact that evolution under antibiotic pressures can select for otherwise less fit proteins [61] and consequently, predictions should not be interpreted as assessments of protein viability or fitness. This is by design, as our goal is to estimate the probability that a variant, once observed in a population, confers resistance and not to speculate on its emergence likelihood. We therefore avoid incorporating general substitution fitness penalties past inherent information from ESM2, as this functionality would also raise biosafety concerns.

It is acknowledged however, that assessing the ability to predict out of distribution is difficult in sparse data settings such as this and much of the *in silico* DMS heatmaps cannot be verified, with the model likely overfitting on the given dataset. That said, the ability to generalize gene information across species, combined with differing hold-out strategy performance, supports the broader validity of this framework. We also recognize that AMR is a multi-layered phenomenon not fully captured by point mutation models, as resistance arises through numerous pathways, including horizontal gene transfer, regulatory changes, and epistasis, which fall outside the scope of this framework. Moreover, the vast theoretical mutational landscape far outstrips the limited catalogue of validated variants, all of which complicates benchmark creation and model training. Our bottom-up approach isolates one tractable component of the problem, yet it must be integrated with orthogonal methods (e.g. efflux pump profiling, promoter region analysis, population level dynamics) to yield a comprehensive risk assessment.

In this work we have focused on delivering a practical tool using risk-based scores, calibration analysis and provision of an easy-to-use pipeline. This emphasis on the assessment of generalised risk, as opposed to metric optimisation on a static benchmark set, is better suited where predictions will be made out-of-distribution and is more in line with clinical decision-making tools. The full training or prediction only workflow can be executed through a single command interface with a user supplied controlling configuration file, facilitating straightforward use.

Moving forward, AMRscope could be extended by integrating more structure-aware and antibiotic-specific features into our embedding, particularly for difficult to predict targets. Additionally, we hope that as public databases continue to collate data on both resistant and sensitive mutations it will provide updated training data, and modular ML pipelines will be expanded to collectively address the multiple layers of AMR evolution. Such integrated platforms will empower real-time genomic surveillance, guiding targeted interventions before resistance becomes widespread.

## Data and Code Availability

The codebase used for model training, evaluation and risk calibration is available at https://github.com/drjjwood/MUTscope and includes all scripts necessary to reproduce the main results of the study. This repository provides:

- Pipeline for protein language model (PLM) embedding, model training, and evaluation
- Notebooks for calibration analysis and DMS visualisations
- Configuration files and environment dependencies for full reproducibility

To support transparency while maintaining clarity, ancillary scripts (e.g., for initial dataset construction, deprecated feature-based models and embeddings) are archived separately.

## Methods

### Ligand Docking

Protein and ligand structures were downloaded from RCSB PBD for PncA (3PL1) [62], KatG (1SJ2) [63], Alr (1XFC) [64], GidB (7CFE), pyrazinamide (PZA), isoniazid (NIZ), cycloserine (DCS) and streptomycin (SRY), and molecular docking performed using Autodock vina (v1.2.5) [29]. Structures and ligands were prepared for docking in PyMOL by removal of water molecules and non-protein atoms, followed by hydrogen addition. The ligand was positioned based on homologous crystal complexes or literature-reported contact residues where available and saved as a cleaned SDF file. Grid box parameters for docking were automatically generated using AutoGrid via the prepare_gpf.pyscript from AutoDockTools, and docking was performed using Vina with AD4 scoring, exhaustiveness set to 32, and two independent runs to assess pose consistency.

The top-ranked poses were evaluated for binding affinity and RMSD relative to the best mode and docking outputs were visualised in PyMOL. Intermolecular interactions were analysed using Arpeggio [65], with attention to van der Waals contacts, polar and hydrophobic interactions, and potential steric clashes. Poses exhibiting strong binding affinity, minimal clash interactions, and multiple stabilizing contacts were prioritised and the best pose used for feature extraction.

### Feature extraction

Features were curated to cover different aspects of mutational impact as follows;

#### Stability

Stability change upon mutation was predicted using DDMut [30] change in ΔΔG and FoldX predicted total energy [31].

#### Evolutionary conservation

ConsurfDB [32] summaries were downloaded for each structure (KatG 1SJ2 chain B, Alr 1XFC Chain B, PncA 3PL1 chain A, RpoB 6DVC chain C, EmbB 7BVF chain B, GidB 7CFE chain A) and parsed to extract per residue conservation scores.

#### Residue properties

Secondary structure (DSSP), accessible surface area (ASA; Shrake-Rupley algorithm), relative solvent accessibility (RSA) and residue depth were extracted using biopython.PDB [34] and residue hydrophobicity calculated using the Kyte-Doolittle scale with a nine amino acid rolling window [37]. Amino acid substitution values were extracted for each mutation for all matrices in amino acid index 2 [33], covering a wide range of property measures.

#### Ligand metrics

Docked structures were uploaded to PLIP [36] and any residues with protein-ligand hydrogen bond distances, hydrophobic interaction distances and salt bridge distances were recorded. Complexes were also used as inputs to mmCSM-lig [35] together with the list of mutations and SMILES codes and resulting protein-ligand affinity changes extracted per mutation. Residue distance to the centre of the docked ligand was also calculated per mutation using biopython.PDB package.

### Feature based modelling

Extracted features were combined into a single dataset, ensuring there were no duplicate mutations, and sensitive and resistant datapoints encoded with 0 (negative class)/1 (positive class) labels. In the case of residues where there were no bond distances to the ligand, missing values were filled with double the maximum recorded bond distance for that bond type. Data was randomly split into 80% training and 20% test set, stratified by labels before min-max scaling was performed on the training set to bring all values into the same range, and this scaler was then applied to the test set. Either LogisticRegression (LR, sklearn), GradientBoostingClassifier (GBC, sklearn), GaussianProcessClassifier (GP, sklearn), or BalancedRandomForestClassifier (BRF, imblearn) were initialised on the full feature set. ‘GridSearchCV’ was used to tune GBC and BRF models across a range of n_estimators (5,50,100,250), min_samples_split (2,4,8,10), min_samples_leaf (1,3,4,5) and max_depths (1,3,7,9,20,40), and the GP over a range of kernels (RBF, DotProduct, Matern, RationalQuadratic, WhiteKernel). The final selected parameters for the combined models were as follows: BRF [min_samples_leaf = 1, min_samples_split = 2, max_depth= 20, n_estimators = 100], GBC [min_samples_leaf = 1, min_samples_split = 8, max_depth= 7, n_estimators = 250], GP [Matern Kernel]. All other parameters were used as the package defaults. Automatic feature selection was performed using ‘SelectFromModel’ and the final set of features used to retrain and tune each model. Feature importance was analysed through direct plotting of model weights or using SHAP summary plots and predictions used to calculate balanced accuracy, F1 score, MCC, Precision and Recall.

### Mutation dataset construction

Data from Tbprofiler, CARD, AMRFinder, PointFinder and Pathogenwatch were downloaded and processed to return AMR associated missense mutations only, wild type protein sequences were obtained from the corresponding databases and mutations applied to generate full mutant sequences [20, 38-40, 42].

To construct a sensitive (negative class) dataset from AllTheBacteria [43], the metadata was filtered to identify sample IDs for organisms of interest with no resistant (positive class) AMRFinder results. For each organism, up to 300 assemblies were downloaded and processed with our variant-calling pipeline. Each assembly was aligned to the organism-specific reference genome using BWA-MEM, and variants were called with bcftools. Gene models for each organism were parsed from GFF3 files, retaining only features annotated as ‘gene’. SNPs were mapped to genes by intersecting VCF positions with gene intervals on matching chromosomes/contigs, accounting for strand orientation. For each SNP, the wild-type gene nucleotide sequence was extracted from the reference FASTA and a mutant sequence was synthesized by substituting the alternate allele at the gene-relative position. Both wild-type and mutant coding sequences were translated to amino acids, and the mutation was classified as synonymous or missense by comparing the affected position. Final per-sample tables were written to CSV for downstream analysis, in some cases down-sampling was applied to the negative set by organism to generate a balanced spread of data.

All the above pre-processed data was combined, any duplicate mutations were dropped and naming conventions of organisms, genes and antibiotics standardised as much as possible. In cases where multiple wild type sequences were found for the same WT variant, these were resolved manually on a case-by-case basis, choosing a single sequence for a single data source but keeping variants across data sources to allow for database variation potentially due to other biological contexts. Finally, the dataset was filtered for the specified suspected direct gene targets, which were identified through manual literature searches.

### Mutation dataset splitting

We implemented multiple independent train/validation/test splitting strategies on the full protein sequences, each holding out approximately 10% of the data as a global test set, followed by five-fold cross-validation on the remaining data.

#### Random stratified splits

Five-fold stratified cross-validation was performed using the label column to preserve the resistant/sensitive class distribution in each fold.

#### Grouped splits by organism, gene, or antibiotic

All samples from a given subgroup (e.g. E. coli, *M. tuberculosis*) were assigned together to either the held-out test set or the train/validation pool, which was further split into five folds using GroupKFold. For antibiotics, grouping was performed first on the resistant subset; the resistant samples were then randomly assigned to test or train/validation to balance class proportions.

#### Homology-aware splits

To minimise data leakage from homologous sequences, full length proteins were clustered using a combined CATH domain architecture and CD-HIT strategy. All unique wild-type protein sequences were (i) queried against the CATH-S35 database [44] (downloaded from the Orengo Lab FTP) using BLASTP (v2.13, E-value ≤ 1×10^−3^) to assign CATH domain architectures, and (ii) clustered at 40% sequence identity using CD-HIT [45] (v4.8; -c 0.4 -n 2) to capture broader homology relationships. Proteins sharing either the same CATH domain architecture (cath_group) or CD-HIT cluster (cluster_id) were connected in a homology graph, and the connected components of this graph defined indivisible homology groups. Entire homology groups were assigned together to the test or train/validation partitions: ∼10% of samples were held out as the global test set using GroupShuffleSplit, and the remaining groups were split into five training/validation folds using GroupKFold.

### PLM modelling

Mutation-level data was collated into a tabular format containing a unique identifier, wild-type sequence, mutant sequence, amino-acid position, wild-type residue, and mutant residue. Each sequence was truncated to a maximum length of 1,020 residues while preserving the mutated position index in frame. For each mutation, both the wild-type and mutant sequences were tokenized and embedded using a protein language model (PLM), of either esm1b_t33_650M_UR50S, esm2_t36_3B_UR50D [17], or SaProt_650M_AF2 [46], extracting the final-layer embedding at the mutated position for both wild-type and mutant sequences. In this case ‘frozen’ weights of pretrained PLMs were used, with no additional pre-training performed before generation of embeddings. These two embeddings were concatenated into a single feature vector, appended with the log-likelihood ratio of the mutant versus wild-type residue (final embedding dimension 5,121) and optionally with additional hand-curated features (Table 1). All appended curated features were min-max scaled prior to training and the final embedding dataloaders created with batch size 16.

The resulting vectors were used as input to a range of classifier heads in, including 1- or 2-layer perceptrons (MLP1lyr/MLP2lyr), support vector classifiers (SVC), gradient boosting classifiers (GBC), and balanced random forests (BRF). Hyperparameter optimization for the selected MLP1lyr model was performed on the training/validation set using random search and logged with Weights & Biases. The search spanned dropout (0.2–0.6), learning rate (10^-8-10^-2), LR decay (0.1–0.5), weight decay (0-10^-2), hidden layer sizes (num_hidden1: 128–1,024; num_hidden2: 64–512), optimizers (Adam/AdamW), schedulers (step or cosine annealing), and training epochs (10–30), with a maximum of 250 runs.

Our final model was a single-layer MLP (MLP1lyr) with ReLU activations and dropout = 0.5. Early stopping was applied with a patience of 20 epochs, and the best checkpoint was selected based on minimal validation binary cross-entropy loss. Training was implemented in PyTorch Lightning with DeepSpeed mixed-precision, acceleration using AdamW as the optimizer (learning rate 1×10^-5, betas= (0.9,0.999), eps 1×10^-8, weight decay1×10−5) with a ReduceLROnPlateau scheduler (factor 0.1, patience 3) under DeepSpeed, or StepLR (step size 5, γ=0.1) otherwise. Class balance was not a major issue in this dataset (positive : negative ∼44:56%), so no reweighting or resampling was applied.

After training, the best model was reloaded for evaluation on the corresponding held-out test set. Performance was assessed using accuracy, precision, recall, F1-score, MCC, and Area Under the Receiver Operating Characteristic Curve (AUROC). For interpretability and downstream visualisation, embeddings were also projected into lower-dimensional spaces using PCA (50 components) and t-SNE, enabling qualitative inspection of separation between biologically relevant groups. As a negative control, the workflow was repeated with shuffled training labels.

The model embedding type, classifier type, hyperparameters, data paths, and logging options are controlled via a single user-specified YAML configuration file.

### Benchmark models

The random test set was used to generate predictions for three comparator models;

#### ESM2

The ‘Facebook/esm2_t36_3B_UR50D’ model to compute Δlog-probability (ΔlogP) for each single amino-acid mutation [17]. The wild-type sequence was masked at the mutated residue, and the log-probability difference between the mutant and wild-type residue at the masked position was used as a mutation score. Negative ΔlogP values indicate less likely mutations under the model.

#### ProtT5

The ‘Rostlab/prot_t5_xl_uniref50’ encoder was used to compute cosine similarity between the wild-type and mutant residue embeddings at the mutated position [48]. A lower similarity score indicates a stronger perturbation.

#### VESPAG

VESPAG was cloned from https://github.com/jschlensok/vespag.git and ran as in the readme to output mutation-level probabilities [47].

Binary predictions from each mode were used to calculate accuracy, Precision, Recall, F1-score and MCC for comparison.

### Calibration and Risk Stratification

We assessed whether the model’s predicted probabilities reflected the true likelihood of a mutation being resistance-associated. For each of the five cross-validation folds, predictions on the held-out test set were grouped into 20 evenly spaced probability bins between 0 and 1. Within each bin, we calculated the fraction of mutations that were truly resistance-associated, which we refer to as the “observed positive rate.” Plotting the observed positive rate against the model’s predicted probability produced a reliability curve, where perfect calibration would follow the diagonal. We also summarised calibration across folds by averaging these curves and shading ±1 standard deviation.

We reported two standard probabilistic metrics: Expected Calibration Error (ECE. – equation 1) and the Brier score (equation 2). ECE measures how far, on average, predicted probabilities deviate from the observed fraction of positives across all bins:

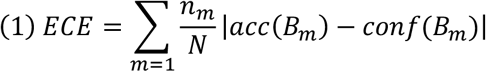

where *B*_*m*_ is a probability bin, *n* _*m*_ is the number of predictions in that bin, *N* is the total number of predictions, *acc*(*B*_*m*_) is the fraction of true positives in that bin, and *con f*(*B*_*m*_) is the mean predicted probability. The Brier score is the mean squared difference between the predicted probability and the true label across all samples:

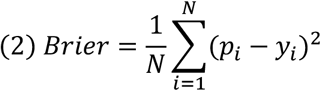

Lower values indicate better probabilistic accuracy.

Finally, we converted predicted probabilities into five risk categories, where probabilities of 0–0.1 were labelled Very Low, 0.1–0.3 Low, 0.3–0.6 Medium, 0.6–0.9 High, and 0.9–1.0 Very High risk. To ensure stable category estimates, we pooled predictions from all training, validation, and test partitions across the five folds when reporting the mean predicted probability and the observed positive rate for each category. These risk bins are for interpretation only and were not used during model training or threshold selection.

## Author Contributions

J.J.W. performed data curation, machine learning model development, analysis and manuscript writing. S.P. contributed guidance on study design and interpretation. D.B.A. provided computational resources and domain-specific guidance. N.F. conceived, designed and supervised the project. All authors contributed to revising the manuscript.

