## Supplementary for "Risk‑Based Prediction of Novel AMR Variants Using Protein Language Models"

**Supplementary Material**

| Gene | **Antibiotic** | **Protein Function** | Ref |
| --- | --- | --- | --- |
| atpE | bedaquiline | ATP synthase subunit C | [1] |
| ddn* | pretomanid and delamanid | deazaflavin-dependent nitroreductase enzyme | [2] |
| EFTu | kirromycin | Elongation Factor | [3] |
| embA | ethambutol | arabinosyltransferases | [4] |
| EmbB |  |  |  |
| embC |  |  |  |
| ethA* | ethionamide | monooxygenase | [5] |
| folA | trimethoprim | dihydrofolate reductase | [6] |
| folC | aminosalicylates | dihydrofolate synthetase | [7] |
| folP | trimethoprim | dihydrofolate reductase | [6] |
| fusA | fusidic acid | elongation factor G | [8] |
| grlA | quinolones and fluoroquinolones | subunit of DNA topoisomerase IV | [9] |
| grlB |  |  |  |
| gyrA |  | subunit of DNA gyrase | [10] |
| gyrB |  |  |  |
| ileS | mupirocin | isoleucyl-tRNA synthetase | [11] |
| inhA | isoniazid | enoyl ACP reductase | [12] |
| katG* |  | catalase-peroxidase | [13] |
| lpxC | lpxC inhibitors | metalloamidase | [14] |
| mshA* | ethionamide | glycosyltransferase | [15] |
| murA | fosfomycin | UDP-N-acetylglucosamine enolpyruvyl transferase | [16] |
| nfsA* | nitrofurantoin | oxygen-insensitive nitroreductases | [17] |
| pbp1 | β-lactam antibiotics | bacterial cell wall synthesis | [18] |
| pbp1a |  |  | [19] |
| pbp2x |  |  | [20] |
| pbp3 |  |  | [21] |
| pncA* | pyrazinamide | pyrazinamide | [22] |
| rplB | bactobolins | L2 ribosomal protein | [23] |
| rplC | linezolid | L3 ribosomal protein | [24] |
| rplD | macrolides | L4 ribosomal protein | [25] |
| rplV |  | L22 ribosomal protein | [26] |
| rpoB | rifampicin | beta subunit of RNA polymerase | [27] |
| rpsA | pyrazinamide | ribosomal protein S1 | [28] |
| rpsE | spectinomycin, streptomycin | ribosomal protein S6 | [29] |
| rpsJ | tetracycline | ribosomal protein S10 | [30] |
| rpsL | streptomycin | ribosomal protein S12 | [31] |

**Table S1. Direct Antibiotics Binding Genes.** List of all potential direct antibiotic targets identified through literature search. (*prodrug activation)

| **organism** | **EFTu** | **atpE** | **ddn** | **embA** | **EmbB** | **embC** | **ethA** | **folA** | **folC** | **folP** | **fusA** | **grlA** | **grlB** | **gyrA** | **gyrB** | **ileS** | **inhA** | **KatG** | **lpxC** | **mshA** | **murA** | **nfsA** | **pbp1** | **pbp1a** | **pbp2x** | **pbp3** | **pbp4** | **PncA** | **rplB** | **rplC** | **rplD** | **rplV** | **RpoB** | **rpsA** | **rpsE** | **rpsJ** | **rpsL** | **Total** |
| --- | --- | --- | --- | --- | --- | --- | --- | --- | --- | --- | --- | --- | --- | --- | --- | --- | --- | --- | --- | --- | --- | --- | --- | --- | --- | --- | --- | --- | --- | --- | --- | --- | --- | --- | --- | --- | --- | --- |
| **Acinetobacter_baumannii** | - | - | - | - | - | - | - | - | - | - | - | 0/4 | - | 0/2 | - | - | - | - | 0/1 | - | - | - | - | - | - | 0/2 | - | - | - | - | - | - | 0/1 | - | - | - | - | 0/10 |
| **Bacillus_subtilis** | - | - | - | - | - | - | - | - | - | - | - | - | - | - | - | - | - | - | - | - | - | - | - | - | - | - | - | - | - | - | - | - | 0/5 | - | 0/1 | - | - | 0/6 |
| **Bartonella_bacilliformis** | - | - | - | - | - | - | - | - | - | - | 2/0 | 3/0 | 1/0 | 7/2 | 2/0 | 4/0 | - | - | - | - | 6/0 | - | - | - | - | - | - | 1/0 | 1/0 | 1/0 | 1/0 | 1/0 | 2/0 | 3/0 | - | - | - | 35/2 |
| **Burkholderia_cepacia** | - | - | - | - | - | - | - | - | - | - | - | - | - | 0/3 | - | - | - | - | - | - | - | - | - | - | - | - | - | - | - | - | - | - | - | - | - | - | - | 0/3 |
| **Burkholderia_pseudomallei** | - | - | - | - | - | - | - | - | - | - | - | - | - | 0/4 | - | - | - | - | - | - | - | - | - | - | - | - | - | - | - | - | - | - | - | - | - | - | - | 0/4 |
| **Campylobacter** | - | - | - | - | - | - | - | - | - | - | - | - | - | 0/11 | - | - | - | - | - | - | - | - | - | - | - | - | - | - | - | - | 0/1 | 0/6 | - | - | - | - | 0/4 | 0/22 |
| **Campylobacter_fetus** | - | - | - | - | - | - | - | - | - | 3/0 | 6/0 | - | - | 15/0 | 13/0 | 44/0 | - | - | 5/0 | - | 7/0 | - | - | - | - | - | - | - | 7/0 | 6/0 | 1/0 | 3/0 | 26/0 | - | - | - | 4/0 | 140/0 |
| **Campylobacter_hyointestinalis** | - | - | - | - | - | - | - | - | - | 14/0 | 15/0 | - | - | 17/0 | 15/0 | 17/0 | - | - | 10/0 | - | 18/0 | - | - | - | - | - | - | - | 14/0 | 15/0 | 13/0 | - | 15/0 | - | 3/0 | 3/0 | 4/0 | 173/0 |
| **Capnocytophaga_gingivalis** | - | - | - | - | - | - | - | - | - | - | - | - | - | 0/1 | - | - | - | - | - | - | - | - | - | - | - | - | - | - | - | - | - | - | - | - | - | - | - | 0/1 |
| **Citrobacter_freundii** | - | - | - | - | - | - | - | - | - | - | - | - | - | 0/2 | - | - | - | - | - | - | - | - | - | - | - | - | - | - | - | - | - | - | - | - | - | - | - | 0/2 |
| **Clostridioides_difficile** | 0/1 | - | - | - | - | - | - | - | - | - | - | - | - | 0/14 | 0/12 | - | - | - | - | - | - | - | - | - | - | - | - | - | - | - | - | - | 3/24 | - | - | - | - | 3/51 |
| **Corynebacterium_diphtheriae** | - | - | - | - | - | - | - | - | - | 7/0 | 15/0 | - | - | 16/4 | 13/0 | 29/0 | - | - | - | 16/0 | 18/0 | - | - | - | - | - | - | - | 6/0 | 8/0 | 11/0 | 4/0 | 0/3 | 14/0 | 2/0 | 5/0 | 2/0 | 166/7 |
| **Cutibacterium_acnes** | - | 12/0 | - | - | - | - | - | - | - | 12/0 | 13/0 | - | - | 15/2 | 13/0 | 16/0 | - | - | - | - | - | - | - | - | - | - | - | - | 8/0 | 12/0 | 14/0 | 10/0 | 16/0 | 12/0 | 12/0 | 4/0 | 4/0 | 173/2 |
| **Enterobacter_cloacae** | - | - | - | - | - | - | - | - | - | - | - | 0/1 | - | 0/6 | - | - | - | - | - | - | - | - | - | - | - | - | - | - | - | - | - | - | - | - | - | - | - | 0/7 |
| **Enterococcus_faecalis** | - | - | - | - | - | - | - | - | - | - | - | 0/5 | - | 0/10 | - | - | - | - | - | - | - | - | - | - | - | - | - | - | - | - | - | - | - | - | - | - | - | 0/15 |
| **Enterococcus_faecium** | 0/2 | - | - | - | - | - | - | - | - | - | - | 0/9 | - | 0/17 | - | - | - | - | - | - | 0/3 | - | - | - | - | - | - | - | - | 0/1 | 0/2 | 0/1 | 0/16 | - | - | 0/9 | - | 0/60 |
| **Erysipelothrix_rhusiopathiae** | - | - | - | - | - | - | - | - | - | - | - | - | - | 0/2 | - | - | - | - | - | - | - | - | - | - | - | - | - | - | - | - | - | - | - | - | - | - | - | 0/2 |
| **Escherichia_coli** | 5/19 | 1/0 | - | - | - | - | - | - | 3/0 | 3/7 | 14/0 | 10/24 | 2/16 | 13/24 | 18/9 | 9/0 | - | 17/0 | 1/0 | - | 1/6 | 4/9 | - | - | - | 8/1 | - | 8/0 | 1/0 | 6/0 | - | 1/0 | 18/18 | 14/0 | 1/0 | - | - | 158/133 |
| **Haemophilus_influenzae** | - | - | - | - | - | - | - | - | - | - | - | - | - | - | - | - | - | - | - | - | - | - | - | - | - | 0/5 | - | - | - | - | - | - | - | - | - | - | - | 0/5 |
| **Haemophilus_parainfluenzae** | - | - | - | - | - | - | - | - | - | - | - | 0/1 | - | 0/1 | - | - | - | - | - | - | - | - | - | - | - | - | - | - | - | - | - | - | - | - | - | - | - | 0/2 |
| **Helicobacter_pylori** | - | - | - | - | - | - | - | - | - | - | - | - | - | 0/11 | 0/1 | - | - | - | - | - | - | - | 0/23 | - | - | 0/5 | - | - | - | - | - | - | 0/29 | - | - | - | - | 0/69 |
| **Klebsiella_oxytoca** | - | - | - | - | - | - | - | - | - | - | - | 0/2 | - | 0/2 | - | - | - | - | - | - | - | - | - | - | - | - | - | - | - | - | - | - | - | - | - | - | - | 0/4 |
| **Klebsiella_pneumoniae** | - | - | - | - | - | - | - | - | - | - | - | 0/2 | - | 0/10 | - | - | - | - | - | - | - | - | - | - | - | 0/1 | - | - | - | - | - | - | - | - | - | 0/1 | - | 0/14 |
| **Morganella_morganii** | - | - | - | - | - | - | - | - | - | - | - | - | - | - | 0/1 | - | - | - | - | - | - | - | - | - | - | - | - | - | - | - | - | - | - | - | - | - | - | 0/1 |
| **Mycobacterium_avium** | - | - | - | - | 19/0 | - | - | - | - | 9/0 | 9/0 | - | - | 10/2 | 11/0 | 21/0 | 11/0 | 13/0 | - | 11/0 | 5/0 | - | - | - | - | - | - | 8/0 | 4/0 | - | 4/0 | 4/0 | - | 5/0 | 15/0 | 2/0 | 2/0 | 163/2 |
| **Mycobacterium_leprae** | - | - | - | - | - | - | - | - | - | 0/14 | 4/0 | - | - | 3/1 | 2/3 | 1/0 | - | - | - | - | - | - | - | - | - | - | - | - | - | - | - | - | 0/2 | - | - | - | - | 10/20 |
| **Mycobacterium_tuberculosis** | - | 10/0 | 10/1 | 11/8 | 14/84 | 10/31 | 9/52 | - | 11/15 | - | - | - | - | 11/20 | 11/22 | - | 10/10 | 11/238 | - | 11/1 | 0/1 | - | - | - | - | - | - | 10/276 | - | 10/1 | - | - | 12/121 | 12/9 | - | - | 10/10 | 183/900 |
| **Mycobacterium_tuberculosis_variant_bovis** | - | - | - | - | 0/3 | - | - | - | - | - | - | - | - | - | - | - | - | - | - | - | - | - | - | - | - | - | - | - | - | - | - | - | - | - | - | - | - | 0/3 |
| **Mycobacteroides_abscessus** | - | 0/2 | - | - | - | - | - | - | - | - | - | - | - | - | - | - | - | - | - | - | - | - | - | - | - | - | - | - | - | - | - | - | - | - | - | - | - | 0/2 |
| **Mycoplasma_genitalium** | - | - | - | - | - | - | - | - | - | - | - | 0/13 | - | 0/2 | - | - | - | - | - | - | - | - | - | - | - | - | - | - | - | - | - | - | - | - | - | - | - | 0/15 |
| **Neisseria_gonorrhoeae** | - | - | - | - | - | - | - | - | 5/0 | 0/3 | - | 0/11 | 0/1 | 0/6 | 0/3 | 12/0 | - | - | - | - | - | - | 0/1 | - | - | - | - | - | - | - | 0/1 | - | 6/5 | - | 0/2 | 0/3 | - | 23/36 |
| **Neisseria_meningitidis** | - | 1/0 | - | - | - | - | - | - | - | - | - | - | - | 2/3 | - | - | - | - | - | - | - | - | - | - | - | - | - | - | 3/0 | - | - | - | 4/6 | - | - | - | - | 10/9 |
| **Pseudomonas_aeruginosa** | 4/0 | - | - | - | - | - | - | - | - | 8/0 | - | 2/2 | 1/6 | 2/12 | 16/9 | 2/0 | - | - | - | - | 1/0 | - | 6/0 | - | - | 4/21 | - | - | 0/2 | 3/0 | 3/0 | - | 3/0 | 5/0 | 1/0 | - | 1/0 | 62/52 |
| **Salmonella** | - | - | - | - | - | - | - | - | - | 0/3 | - | 0/8 | 0/8 | 0/24 | 0/7 | - | - | - | - | - | - | - | - | - | - | - | - | - | - | - | - | - | - | - | - | - | - | 0/50 |
| **Salmonella_enterica** | - | - | - | - | - | - | - | - | - | - | - | 70/1 | 27/0 | - | - | - | - | - | - | - | 18/0 | - | - | - | - | - | - | 9/0 | - | 5/0 | 4/0 | 1/0 | - | - | - | - | - | 134/1 |
| **Salmonella_typhi** | - | - | - | - | - | - | - | - | - | - | - | 0/3 | 0/2 | 0/7 | 0/2 | - | - | - | - | - | - | - | - | - | - | - | - | - | - | - | - | - | - | - | - | - | - | 0/14 |
| **Serratia_marcescens** | - | - | - | - | - | - | - | - | - | - | - | - | - | 0/3 | - | - | - | - | - | - | - | - | - | - | - | - | - | - | - | - | - | - | - | - | - | - | - | 0/3 |
| **Shigella_flexneri** | - | - | - | - | - | - | - | - | - | - | - | - | - | 0/2 | - | - | - | - | - | - | - | - | - | - | - | - | - | - | - | - | - | - | - | - | - | - | - | 0/2 |
| **Staphylococcus_aureus** | - | - | - | - | - | - | - | - | - | 0/5 | 0/68 | 0/14 | 0/16 | 0/12 | 0/12 | 0/6 | - | - | - | - | 0/7 | - | - | - | - | - | 0/4 | - | 1/0 | 1/4 | 2/1 | 1/0 | 3/36 | - | - | 0/1 | 1/0 | 9/186 |
| **Staphylococcus_pseudintermedius** | - | - | - | - | - | - | - | - | - | 5/0 | 2/0 | 7/6 | 7/0 | 13/6 | 9/0 | 12/0 | - | - | - | - | 4/0 | 10/0 | - | - | - | - | 10/0 | - | 2/0 | - | - | 1/0 | 1/2 | 7/0 | - | - | - | 90/14 |
| **Streptococcus_agalactiae** | - | - | - | - | - | - | - | - | - | - | - | - | - | - | - | - | - | - | - | - | - | - | - | - | 0/1 | - | - | - | - | - | - | - | - | - | - | - | - | 0/1 |
| **Streptococcus_pneumoniae** | - | - | - | - | - | - | - | 0/1 | - | 20/0 | 11/0 | 12/10 | 12/4 | 12/6 | 13/1 | 17/0 | - | - | - | - | 8/0 | - | - | 13/2 | 15/6 | 8/0 | - | - | 4/0 | 6/0 | 3/0 | 1/0 | 9/0 | 11/0 | - | 1/0 | - | 176/30 |
| **Streptococcus_pyogenes** | - | - | - | - | - | - | - | - | - | 11/1 | 8/0 | 17/0 | 16/0 | 10/0 | 11/0 | 15/0 | - | - | - | - | 11/0 | - | - | 12/0 | 10/3 | 11/0 | - | - | 6/0 | 6/0 | 6/0 | 4/0 | 10/0 | 10/0 | 3/0 | 1/0 | 1/0 | 179/4 |
| **Ureaplasma_urealyticum** | - | - | - | - | - | - | - | - | - | - | - | 0/4 | - | - | 0/2 | - | - | - | - | - | - | - | - | - | - | - | - | - | - | - | - | - | - | - | - | - | - | 0/6 |
| **Vibrio_cholerae** | - | 1/0 | - | - | - | - | - | 10/0 | 11/0 | 10/0 | 11/0 | - | 5/2 | 18/2 | 10/0 | 13/0 | - | 10/0 | 2/0 | - | 10/0 | 13/0 | - | - | - | - | - | - | 5/0 | 1/0 | - | - | 13/0 | 1/0 | - | - | 3/0 | 147/4 |
| **Vibrio_parahaemolyticus** | - | - | - | - | - | - | - | - | - | - | - | 0/2 | - | 0/2 | - | - | - | - | - | - | - | - | - | - | - | - | - | - | - | - | - | - | - | - | - | - | - | 0/4 |
| **Vibrio_vulnificus** | - | - | - | - | - | - | - | 7/0 | 1/0 | 2/0 | 2/0 | 9/1 | 6/0 | 11/2 | 27/0 | 10/0 | - | 5/0 | 3/0 | - | 4/0 | - | - | - | - | - | - | - | - | - | 1/0 | - | 1/1 | - | - | - | - | 89/4 |
| **Total** | 9/22 | 25/2 | 10/1 | 11/8 | 33/87 | 10/31 | 9/52 | 17/1 | 31/15 | 104/33 | 112/68 | 130/123 | 77/55 | 175/240 | 184/84 | 222/6 | 21/10 | 56/238 | 21/1 | 38/1 | 111/17 | 27/9 | 6/24 | 25/2 | 25/10 | 31/35 | 10/4 | 36/276 | 62/2 | 80/6 | 63/5 | 31/7 | 142/269 | 94/9 | 37/3 | 16/14 | 32/14 | 2123/1784 |

**Table S2. Full dataset by organism and gene.** Count of all datapoints for every gene and species, in the format sensitive/resistance, with column and totals for each group. All genes have both sensitive and resistant representation.

| **Set number** | **group** | **Hold out test groups** | **Total test points**  **(n resistant/ n sensitive)** | **Percentage resistant** |
| --- | --- | --- | --- | --- |
| **1** | organism | 'Burkholderia_cepacia', 'Campylobacter_fetus', 'Capnocytophaga_gingivalis', 'Citrobacter_freundii', 'Mycobacterium_leprae' | 176 (26/150) | 14.8 |
| **2** |  | 'Mycobacterium_avium', 'Mycobacterium_tuberculosis', 'Mycobacterium_tuberculosis_variant_bovis', 'Staphylococcus_pseudintermedius', 'Ureaplasma_urealyticum' | 1506 (925/581) | 61.4 |
| **3** |  | 'Cutibacterium_acnes', 'Erysipelothrix_rhusiopathiae', 'Morganella_morganii', 'Salmonella_typhi', 'Streptococcus_pyogenes' | 375 (23/352) | 6.1 |
| **4** |  | 'Burkholderia_pseudomallei', 'Enterococcus_faecium', 'Pseudomonas_aeruginosa', 'Staphylococcus_aureus', 'Vibrio_parahaemolyticus' | 373 (302/71) | 81.0 |
| **5** |  | 'Escherichia_coli', 'Haemophilus_influenzae', 'Neisseria_gonorrhoeae', 'Streptococcus_agalactiae', 'Vibrio_cholerae' | 507 (179/328) | 35.3 |
| **1** | homology | 'Cluster 0', 'Cluster 24', 'Cluster 43', 'Cluster 45', 'Cluster 5', 'Cluster 50', 'Cluster 69' | 611 (238/373) | 39.0 |
| **2** |  | 'Cluster 25', 'Cluster 27', 'Cluster 34', 'Cluster 4', 'Cluster 42', 'Cluster 70', 'Cluster 71' | 214 (58/156) | 27.1 |
| **3** |  | 'Cluster 17', 'Cluster 37', 'Cluster 54', 'Cluster 62', 'Cluster 66', 'Cluster 68', 'Cluster 7' | 224 (35/189) | 15.6 |
| **4** |  | 'Cluster 1', 'Cluster 15', 'Cluster 26', 'Cluster 31', 'Cluster 36', 'Cluster 47', 'Cluster 52' | 451 (150/301) | 33.3 |
| **5** |  | 'Cluster 13', 'Cluster 28', 'Cluster 35', 'Cluster 38', 'Cluster 49', 'Cluster 55', 'Cluster 56' | 311 (114/197) | 36.7 |
| **1** | gene | 'EFTu', 'mshA', 'rpsA', 'rpsL' | 248 (46/202) | 18.5 |
| **2** |  | 'embA', 'rplC', 'rplV', 'rpsJ' | 196 (35/161) | 17.9 |
| **3** |  | 'embB', 'lpxC', 'pbp1', 'pncA' | 496 (388/108) | 78.2 |
| **4** |  | 'folA', 'katG', 'rplB', 'rpoB' | 826 (510/316) | 61.7 |
| **5** |  | 'atpE', 'ddn', 'folP', 'ileS' | 421 (42/379) | 10.0 |
| **1** | antibiotic | ‘AMX’ | 72 (36/36) | 50.0 |
| **2** |  | ‘EMB’ | 201 (124/77) | 61.7 |
| **3** |  | ‘INH' | 422 (269/153) | 63.7 |
| **4** |  | ‘PZA' | 434 (285/149) | 65.7 |
| **5** |  | ‘RIF’ | 432 (226/206) | 52.3 |

**Table S3 Summary of hold-out test set splits.** For each non-random hold-out strategy (homology, gene, organism, antibiotic) five independent splits were generated for training and testing and the test set features summarized in the table.

| **Organism** | **Gene** | **Dataset Mutations** | **Antibiotic Resistance** |
| --- | --- | --- | --- |
| *M. tuberculosis.* | RpoB | V146F, V170F, A286V, T400A, E423A, F424L, F424V, T427I, T427P, T427S, T427G, T427A, T427N, S428G, S428I, S428R, S428T, Q429K, Q429H, Q429P, Q429L, L430P, L430R, L430V, S431G, S431R, Q432P, Q432E, Q432L, Q432K, Q432H, M434L, M434T, M434V, M434I, D435A, D435N, D435G, D435E, D435H, D435Y, D435F, D435V, D435L, Q436L, Q436N, Q436P, N437D, N437I, N437Y, N437H, N437S, P439S, P439L, S441L, S441A, S441V, S441M, S441W, S441Q, G442E, G442W, L443F, L443W, T444S, T444I, T444P, H445Y, H445C, H445D, H445T, H445P, H445L, H445F, H445S, H445G, H445R, H445Q, H445N, K446T, K446E, K446Q, K446R, R448K, R448Q, L449M, S450W, S450G, S450L, S450Q, S450V, S450A, S450F, S450Y, S450M, S450C, A451G, A451V, L452P, L452V, L452M, P454R, P454L, P454H, P454S, E460G, I480T, I480V, P483L, I491F, S493L, F505L, T508N, T508H, T508S, T508P, T508A, R552C, E592D, E761D, G981D | rifampicin |
| *E. coli* | GyrB | R136S, R136H, R136E, R136C, R136G, R136L, R136I, | aminocoumarin |
|  |  | D426N, K447E | quinolone |
| *Salmonella* | GrlA | T57S, T66I, G78D, S80I, S80R, E84G, E84K, W106G | quinolone |
| *P. aeruginosa* | PBP3 | G63S, Y367C, S368L, H394R, A419G, N427S, N427L, L434V, A454V, Q458R, L461V, V471G, Q475R, R504L, R504C, F507L, V523A, V523M, P527S, F533L, S538L | carbapenem |

**Table S4. Dataset mutations for in silico predictions.** Mutations present in our initial dataset for the genes investigated by in silico DMS.

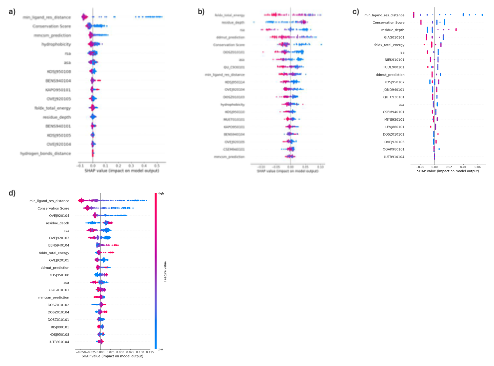

**Figure S1 Feature importance of M. tuberculosis single gene models.** SHAP importance of the features used in balanced random forest models for a) RpoB b) PncA c) KatG d) EmbB.

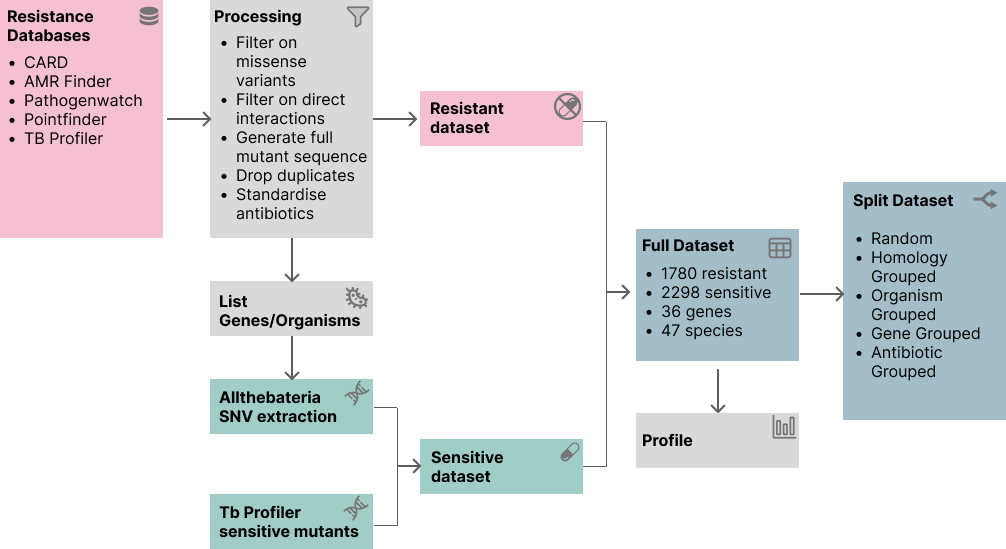

**Figure S2 Data collection and processing workflow.** Data was downloaded from the Comprehensive Antibiotic Resistance Database (CARD), AMR finder, pathogen watch and pointfinder databases and filtered to get missense variants only. This data was combined with the *M. tuberculosis* dataset and processed to remove duplicates, to standardise antibiotics and filter for genes which are known or suspected antibiotic interaction partners. An additional negative dataset for complementary genes was constructed from Allthebacteria to generate a final dataset which was subjected to splitting with five different hold-out strategies.

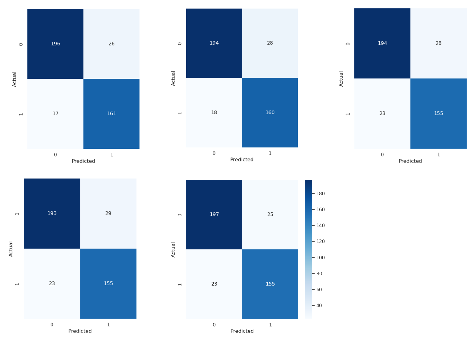

**Figure S3 Confusion matrices of each cross-validation fold.** Five confusion matrices from for the hold-out test set for models generated from five-fold cross validation.

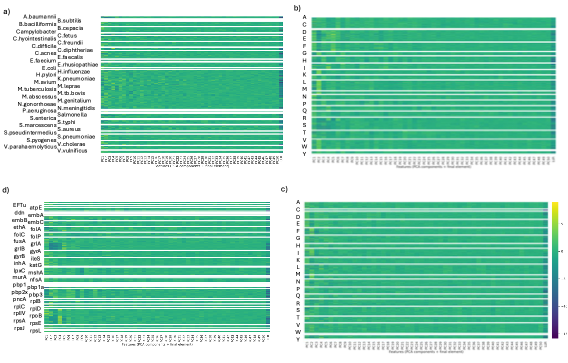

**Figure S4. Dimensionality reduction of embeddings.** Heatmap of PCA reduced (50 components) input vectors for an illustrative validation set split by a) WT amino acid b) mutant amino acid c) gene d) organism.
